# Genome-wide conservation and locus-specific coding variation in *Teratoramularia rumicicola*, a candidate microbial herbicide for *Rumex* weeds

**DOI:** 10.64898/2026.10.17.759301

**Authors:** Masataka Izumi, Tomomasa Oka

## Abstract

The *Rumex* pathogen *Teratoramularia rumicicola* is a candidate microbial herbicide, but its functional genomic repertoire and variation among isolates remain poorly characterised. We generated a PacBio HiFi reference and evidence-supported gene annotation for TR4 and a short-read assembly for GR1, then compared the isolates using assembly- and read-based analyses. The TR4 and GR1 nuclear assemblies comprised 22.34 Mb (13 contigs; N50, 2.90 Mb) and 21.23 Mb (486 contigs; N50, 0.350 Mb), respectively. The TR4 annotation predicted 8,898 protein -coding genes, and the representative protein set was 96.7% complete in the Benchmarking Universal Single-Copy Orthologues assessment. Functional annotation identified 593 putatively secreted proteins, 258 candidate effectors, 355 carbohydrate -active enzyme (CAZyme) candidates and 20 predicted secondary-metabolite biosynthetic regions. Seven secreted CAZymes were prioritised as potential host-barrier-associated factors based on predicted activities and homology to virulence -related entries in the Pathogen–Host Interactions database. Despite 99.81% average nucleotide identity across aligned regions, read-based comparison identified 28,040 high-confidence isolate-fixed differences. Of 158 reconstructed candidate GR1 alleles, 147 showed strong assembly support. Notable differences included predicted truncations of two Zn(II)₂Cys₆ transcription factors, one adjacent to an intact nonribosomal peptide synthetase-like core gene, and a separate gene showing a localised repeat-induced point mutation (RIP)-like substitution pattern and predicted premature termination. Together, these resources and candidate loci provide a foundation for investigating pathogenicity and strain variation relevant to microbial herbicide development.

## 1. Introduction

Perennial docks, particularly *Rumex crispus* L. and *Rumex obtusifolius* L., are troublesome weeds in temperate grasslands and pastures. Their persistent root systems, capacity for regrowth and prolific seed production promote long -lived infestations that can reduce forage yield and quality (Zaller, 2004; Klötzli et al., 2024). Selective chemical control is difficult in mixed grass–legume swards because herbicides targeting broad-leaved weeds may also injure desirable forage legumes (McCurdy et al., 2013). Asulam has long been used for dock control in grasslands (Brock, 1972), but asulam-sodium is not approved in the European Union (European Commission, 2024). These constraints create a need for additional selective approaches to *Rumex* management, including fungal biological control.

Fungal biological control of *Rumex* weeds has previously been investigated using *Uromyces rumicis* and *Ramularia rubella*, although practical application remains limited (Schubiger et al., 1986; Huber-Meinicke et al., 1989; Zaller, 2004). *Teratoramularia rumicicola* is a ramularioid foliar pathogen that has been reported from *Rumex* hosts in Japan and Korea (Videira et al., 2016; Choi et al., 2025; Izumi and Sato, 2025). Strains TR4 and GR1 are pathogenic to multiple *Rumex* species, and preliminary evidence indicates TR4 selectivity between target weeds and tested forage species (Izumi and Sato, 2025, 2026). Both strains can also produce conidia in liquid culture, supporting further evaluation of *T. rumicicola* for selective *Rumex* control.

Genome-enabled studies of fungal plant pathogens have identified secreted proteins, candidate effectors, carbohydrate-active enzymes (CAZymes) and secondary-metabolite biosynthetic gene clusters as useful functional repertoires for investigating host interactions and pathogenicity (Kubicek et al., 2014; Lo Presti et al., 2015; Keller, 2019). In the fungal microbial herbicide candidate *Trichoderma polysporum* HZ-31, genome annotation identified putative pathogenicity-related genes, while transcriptome profiling was used to investigate gene expression during infection of *Avena fatua* L. (He and Zhu, 2024; Zhu and He, 2024). Comparative genomic analyses provide additional insights by revealing genomic conservation and isolate-associated sequence variation, which can inform the prioritisation of candidate loci for subsequent functional investigation. For example, combined pangenomic and transcriptomic analyses of *Zymoseptoria tritici* demonstrated the utility of gene conservation and expression data for prioritising candidates involved in virulence (Chen et al., 2023).

To date, no studies on the molecular mechanisms underlying pathogenicity in *T. rumicicola* or comparative genomic analyses between isolates have been reported. The only published genome resource for this species is an unannotated short-read draft assembly of TR4 (Izumi et al., 2026). Strains TR4 and GR1 were isolated from different *Rumex* hosts at geographically distant locations nearly three decades apart (Izumi and Sato, 2025, 2026). These distinct isolation backgrounds provide an opportunity to examine genome-wide conservation and isolate-associated sequence variation.

Here, we generated a highly contiguous PacBio HiFi assembly of TR4 with evidence-supported gene annotation and a short-read genome assembly of GR1. We systematically characterized predicted secreted proteins, candidate effectors, CAZymes and secondary-metabolite biosynthetic regions to establish a molecular catalogue of potential host-interaction factors in *T. rumicicola*. We then compared the isolates using complementary assembly- and read-based approaches to assess genome-wide conservation and identify localized isolate-associated variation with potential coding consequences.

## 2. Materials and Methods

### 2.1 Fungal strains and culture conditions

Two strains of *T. rumicicola* that had been isolated and deposited prior to the present study were examined. Strain TR4 had been obtained by single-conidium isolation from a symptomatic *R. crispus* leaf collected in Tsukuba, Ibaraki Prefecture, Japan, in November 2024, and deposited in the NARO Genebank as MAFF 248116 (Izumi and Sato, 2025). Strain GR1 had been isolated from a naturally infected *Rumex japonicus* Houtt. leaf collected on Hahajima Island, Ogasawara Islands, Tokyo, Japan, in June 1996, and deposited as MAFF 238946 (Izumi and Sato, 2026).

For genomic DNA preparation, TR4 and GR1 were cultured on potato dextrose agar at 25°C in darkness for 27 and 21 days, respectively. Mycelium was scraped from the colony surface without underlying agar and stored at −80°C. For RNA preparation, TR4 was cultured in 0.5% yeast extract–malt extract broth as described previously (Izumi and Sato, 2025). A 5-mm mycelial plug was precultured in 50 mL broth at 25°C for 3 days on a rotary shaker (130 rpm); 1% (v/v) was then transferred to 150 mL fresh broth and cultured at 25°C for 7 days on a reciprocating shaker (130 strokes min⁻¹). Biomass was harvested, washed three times with sterile water, flash-frozen in liquid nitrogen and stored at −80°C.

### 2.2 Genomic DNA extraction and sequencing

All genomic DNA and total RNA extraction, library preparation and sequencing were performed by Bioengineering Lab. Co., Ltd. (Sagamihara, Japan). High -molecular-weight TR4 DNA was extracted using a QIAGEN Genomic-tip 20/G column. Short fragments were depleted with Short Read Eliminator XS, and the DNA was sheared to approximately 7–20 kb. A PCR-amplified SMRTbell library was prepared from 20 ng of DNA using SMRTbell Prep Kit 3.0 and Twist Universal Adapter System (10 PCR cycles) and sequenced on a PacBio Revio system. Circular consensus reads were generated using SMRT Link v25.2.0.266456; reads with mean quality below 20 were discarded, and adapters and duplicate reads were removed using lima v2.12.0 and pbmarkdup v1.0.3, respectively.

GR1 DNA was extracted using the MPure-12 system and MPure Bacterial DNA Extraction Kit. A library was prepared from 50 ng of DNA using MGIEasy Fast FS Library Prep Set V2.0 and sequenced on a DNBSEQ-T7 platform to generate 150-bp paired-end reads.

### 2.3 Genome assembly, curation and quality assessment

TR4 HiFi reads were assessed with NanoPlot v1.46.2 (De Coster et al., 2018) and assembled using hifiasm v0.25.0 -r726 (Cheng et al., 2021) with haplotig purging disabled (-l 0). GR1 reads were assessed with FastQC v0.11.9, adapter-trimmed and quality-filtered with fastp v0.20.1 using manually specified adapters and otherwise default settings (Chen et al., 2018), and assembled using SPAdes v4.2.0 in isolate mode (Bankevich et al., 2012).

Putative mitochondrial sequences were removed from both assemblies. For TR4, contigs with GC content below 40% were compared by all -versus-all BLASTN (Camacho et al., 2009). Ten long representatives were confirmed as fungal mitochondrial sequences against the NCBI core nucleotide database; candidates with ≥99% identity and ≥99% query coverage to these representatives were removed. For GR1, a 34,591-bp low-GC, high-coverage mitochondrial contig and five short contigs with nearly full-length similarity were removed; contigs with only short local matches were retained. Contigs shorter than 200 bp were excluded for GenBank submission; subsequent NCBI Foreign Contamination Screen excluded 25 additional short GR1 contigs, yielding the final GR1 nuclear assembly.

The curated TR4 and GR1 nuclear assemblies and the previously published TR4 short-read assembly (Izumi et al., 2026) were evaluated using QUAST v5.3.0 (Gurevich et al., 2013). Completeness was assessed using Benchmarking Universal Single-Copy Orthologues (BUSCO) v6.1.0 (Manni et al., 2021) in genome mode with MetaEuk (Levy Karin et al., 2020) against dothideomycetes_odb12.2. Telomeric repeats were screened by searching the terminal 100 bp of contigs ≥500 bp for at least three consecutive exact copies of 5′-TTAGGG-3′ or 5′-CCCTAA-3′.

### 2.4 RNA extraction, sequencing and read alignment for TR4

Total RNA was extracted from frozen TR4 biomass using the RNeasy Mini Kit with QIAcube Connect. An RNA sequencing (RNA-seq) library was prepared from 300 ng of RNA using the MGIEasy Fast RNA Library Prep Set and sequenced on a DNBSEQ-T7 platform (150-bp paired-end). Reads were assessed using FastQC v0.11.9 and aligned to the curated TR4 nuclear genome using HISAT2 v2.2.1 with --dta (Kim et al., 2019). Alignments were sorted and indexed using SAMtools (Danecek et al., 2021) and supplied to BRAKER as transcript evidence. The data were not used for quantitative expression analysis.

### 2.5 Repeat masking and structural gene prediction

A de novo repeat library was generated from the curated TR4 assembly using RepeatModeler v2.0.8 with long terminal repeat structural analysis (Flynn et al., 2020) and RMBlast v2.17.1+. The unclassified library was used for soft masking with RepeatMasker v4.2.4 (Smit et al., 2013 –2015). BRAKER v3.0.8 in ETP mode (Gabriel et al., 2024) predicted gene models using the HISAT2 -aligned RNA-seq BAM file and OrthoDB12 Fungi proteins as evidence (Tegenfeldt et al., 2025).

### 2.6 Representative protein set and annotation assessment

One protein-coding transcript model per predicted gene was selected for downstream analyses: .t1 except for g8894, where the structurally complete .t2 replaced a .t1 lacking an annotated start codon at a contig boundary. All selected models contained annotated start and stop codons without internal stops. The representative protein set was assessed using BUSCO v6.1.0 in protein mode against dothideomycetes_odb12.2.

### 2.7 Functional annotation of predicted proteins

The representative TR4 protein set was annotated using eggNOG-mapper v2.1.7 (Cantalapiedra et al., 2021) with DIAMOND v2.2.4 (Buchfink et al., 2021) against eggNOG v5.0.2. After removal of terminal stop symbols, InterProScan v5.78 -109.0 (Jones et al., 2014) was run with InterPro release 109.0. Protein-level annotations are provided in Table S1.

### 2.8 Functional repertoire prediction and PHI-base annotation

Signal peptides were predicted from the representative TR4 protein set using SignalP 6.0 (Teufel et al., 2022) with the organism group set to Eukarya. SignalP -positive proteins were trimmed at the predicted cleavage site and analysed using DeepTMHMM 1.0 (Hallgren et al., 2022). Proteins with predicted transmembrane regions were excluded, and the remainder defined as the putative secretome. Candidate effectors were predicted from the trimmed sequences using EffectorP 3.0 (Sperschneider and Dodds, 2022), retaining apoplastic, cytoplasmic and dual-localized effectors.

CAZymes were annotated using dbCAN3 (Zheng et al., 2023) with HMMdb v14.0. HMMER searches against dbCAN and dbCAN-sub used an E-value threshold of 1 × 10^−15^ and coverage >0.35, whereas DIAMOND searches against CAZy used an E-value threshold of 1 × 10^−102^. Proteins supported by at least two of the three methods were retained.

Biosynthetic gene clusters were predicted using the fungiSMASH web server, based on antiSMASH v8.0.4 (Blin et al., 2025), with the curated TR4 assembly and a GFF3 containing one selected representative transcript model per gene. Detection strictness was set to relaxed; all other settings were defaults.

The representative TR4 protein set was searched against the Pathogen –Host Interactions database (PHI-base) v4.19 (Urban et al., 2025) using DIAMOND v2.2.5 in sensitive mode, with an initial E-value threshold of 1 × 10^−5^ and up to 25 targets per query. The highest-bit-score hit meeting amino-acid identity ≥30%, query and subject coverage ≥60% and E-value ≤1 × 10^−10^ was retained. Hits were classified as pathogenicity-related when the PHI-base phenotype included loss of pathogenicity, reduced virulence, increased virulence or effector activity. For the targeted secreted CAZyme analysis, virulence-related hits were defined as those associated with loss of pathogenicity, reduced virulence or increased virulence, excluding hits associated exclusively with effector activity. These matches were treated as homology evidence rather than evidence of equivalent function in *T. rumicicola*.

### 2.9 PHI-base-guided functional-category enrichment analysis

Functional-category enrichment was evaluated for the putative secretome, candidate effectors, CAZymes, secreted CAZymes, fungiSMASH biosynthetic-region members, core biosynthetic genes and proteins classified as both candidate effectors and CAZymes. One-sided Fisher’s exact tests were used to test for overrepresentation of each category among pathogenicity-related proteins relative to other representative proteins. P-values were adjusted using the Benjamini–Hochberg procedure across the seven categories within each analysis, with adjusted P-values < 0.05 considered significant. The analysis was repeated within proteins having strict PHI-base hits. In a sensitivity analysis, proteins whose only qualifying PHI-base phenotype was effector activity were reassigned from the pathogenicity-related group to the comparison group within the strict-hit background. Statistical analyses were performed in R v4.4.3.

### 2.10 Whole-genome assembly comparison

The curated TR4 HiFi, published TR4 short-read and GR1 short-read assemblies were compared pairwise using dnadiff v1.3 in MUMmer4 v4.0.1 ( Marçais et al., 2018). One-to-one average nucleotide identities and aligned sequence lengths were obtained from dnadiff, and alignment fractions were calculated relative to the respective nuclear assembly spans.

### 2.11 Read mapping and isolate-fixed variant detection

Quality-filtered paired-end reads from the previously published TR4 dataset (Izumi et al., 2026) and the newly generated GR1 dataset were independently mapped to the curated TR4 HiFi nuclear genome using BWA-MEM2 v2.2.1 (Vasimuddin et al., 2019). Alignments were sorted and indexed using SAMtools v1.24, and depth and coverage were assessed using SAMtools and mosdepth v0.3.14 (Pedersen and Quinlan, 2018).

Variants were jointly called using BCFtools v1.24 (Danecek et al., 2021). bcftools mpileup was run with minimum mapping and base qualities of 30 and 20, respectively, a maximum per-sample depth of 10,000, and allelic depth and total depth recorded for each sample. The multiallelic caller was used with both isolates treated as haploid and population-based genotype priors disabled. Variant records were normalized against the TR4 reference, and multiallelic records were split. Biallelic single-nucleotide polymorphisms (SNPs) and short insertions or deletions were retained when site quality was ≥30, read depth was 10 –500 in both isolates, and alternative-allele frequencies were ≤0.10 in TR4 and ≥0.90 in GR1. Alternative-allele frequencies were calculated from allelic read depths. The upper depth threshold was used to exclude unusually high-coverage loci potentially associated with repetitive or collapsed genomic regions. Sites satisfying these within -isolate thresholds are termed isolate-fixed differences here; the term does not imply fixation across the species. Variant positions and predicted effects were annotated using a custom SnpEff database constructed from the representative TR4 gene models (Cingolani et al., 2012).

### 2.12 Candidate-gene selection and allele reconstruction

Genes were selected for allele-level characterization if they contained at least one isolate-fixed SnpEff HIGH-impact variant or ranked in the upper 1% of HIGH- and MODERATE-impact variant density per kilobase of representative coding sequence (CDS). Density ranking was restricted to genes with at least one HIGH- or MODERATE-impact variant and a representative CDS length ≥300 bp.

For each candidate, isolate-fixed variants overlapping the representative TR4 CDS were applied simultaneously to reconstruct the corresponding GR1 coding allele. TR4 and reconstructed GR1 CDSs were compared for length changes, reading -frame preservation, start-codon retention and terminal stop-codon retention. The translated sequences were examined for premature termination and differences in predicted protein length. When the reconstructed GR1 CDS lacked an in-frame stop codon, downstream TR4 genomic sequence was modified with GR1 isolate-fixed variants and searched for an in-frame stop codon within 10 kb in the direction of transcription. CDS reconstruction assumed the TR4 exon–intron structure; variants outside the annotated CDS were excluded from the CDS itself, and potential splice changes were not evaluated. Functional annotations were integrated with reconstructed sequence properties (Table S2). PHI-base similarity was not used for candidate selection.

Separately, seven virulence-related secreted CAZyme genes were exploratorily prioritized as host-barrier-associated candidates based on sequence annotations indicating potential activities related to xylan or hemicellulose, pectin or cuticle modification, and their GR1 alleles were reconstructed using the same procedure. They were not added to the systematic candidate set unless they independently met the predefined variant-based criteria. For locus-level comparison of the g5338–g5339 region, the adjacent core biosynthetic gene g5339 was also reconstructed using the same procedure. Amino-acid substitutions were additionally examined for the seven prioritized CAZyme genes and g5339. Because g8540 contained an N-terminal R3L substitution, its reconstructed GR1 protein was additionally assessed using SignalP 6.0 and EffectorP 3.0 as described in Section 2.8.

### 2.13 Assembly-based evaluation of reconstructed alleles against the GR1 de novo assembly

Reconstructed GR1 CDS sequences from both the systematic candidate set and the targeted seven-gene analysis were aligned to the separately assembled GR1 nuclear genome using minimap2 v2.24 -r1122 in spliced-alignment mode with the splice:hq preset. Primary alignments were generated with secondary alignments suppressed, and the analysis was repeated allowing up to 10 secondary alignments to identify alternative genomic matches.

Alignment support was evaluated using query coverage, mapping quality, alignment multiplicity and sequence agreement calculated as 100 × (1 − de), where de is the gap-compressed divergence reported by minimap2. Alignments were classified as strongly supported when query coverage and sequence agreement were both ≥99.9%, mapping quality was 60 and only one alignment was reported; other alignments were categorized as showing partial coverage, sequence differences or ambiguous mapping.

### 2.14 Locus-specific read and substitution-pattern analyses

The g8812 locus was examined using the TR4 -reference read alignments generated in Section 2.11. Local read depth, mapping quality and paired-read support were assessed to evaluate the reliability of the isolate-fixed variants. The g8812 coding region (ptg000009l:218,884 –219,444) was compared with a nearby RepeatMasker-annotated multicopy repeat (212,101 –213,966), the intervening genomic interval (213,967–218,883) and a higher-coordinate flanking region (219,445 –220,000). C→T and G→A substitutions were combined as C:G-to-T:A changes. Variant densities were calculated using only positions with read depth of 10 –500 in both isolates. CpA/TpG target sites were enumerated from the TR4 reference sequence, and the proportion carrying the corresponding C:G-to-T:A isolate-fixed substitution was calculated for each region.

## 3. Results

### 3.1 Genome assembly and quality assessment of strains TR4 and GR1

After removal of putative mitochondrial sequences, the curated TR4 HiFi nuclear assembly comprised 13 contigs totalling 22.34 Mb, with an N50 of 2.90 Mb (Table 1). Compared with the published TR4 short-read assembly, the number of contigs ≥500 bp decreased from 316 to 13 and the N50 increased from 0.284 to 2.90 Mb. The two TR4 assemblies were nearly identical at the nucleotide level (99.9950% average nucleotide identity). Canonical telomeric repeats were detected at 7 of the 26 HiFi contig ends, consistent with a highly contiguous but not chromosome -complete assembly.

**Table 1.** Assembly statistics and genome completeness of *Teratoramularia rumicicola* strains TR4 and GR1.

| Assembly | Sequencing approach | Contigs $\geq 500$ bp | Assembly span in contigs $\geq 500$ bp (Mb) | Contig N50 (Mb) | Telomeric ends detected/examined | Complete BUSCOs (%) |
| --- | --- | --- | --- | --- | --- | --- |
| TR4 HiFi | PacBio HiFi | 13 | 22.34 | 2.90 | 7/26 | 93.6 |
| TR4 short-read assembly<sup>1</sup> | DNBSEQ-T7 PE150 | 316 | 21.32 | 0.284 | 1/632 | 93.7 |
| GR1 short-read assembly | DNBSEQ-T7 PE150 | 250 | 21.16 | 0.350 | 1/500 | 93.8 |

Contig counts, assembly spans and telomeric-end counts are reported for contigs $\geq 500$ bp. Genome completeness was assessed using BUSCO v6.1.0 in genome mode with MetaEuk against the dothideomycetes\_odb12.2 lineage dataset. Telomeric ends were defined by the presence of at least three consecutive copies of 5'-TTAGGG-3' or 5'-CCCTAA-3' within the terminal 100 bp of contigs $\geq 500$ bp.
<sup>1</sup> Previously published TR4 short-read assembly (Izumi et al., 2026).

The curated GR1 nuclear assembly comprised 486 contigs totalling 21.23 Mb, with an N50 of 0.350 Mb. Of these, 250 contigs were ≥500 bp, accounting for 21.16 Mb (Table 1). BUSCO completeness was similar across the TR4 HiFi, published TR4 short-read and GR1 assemblies (93.6 –93.8%), despite their marked differences in contiguity.

### 3.2 Evidence-supported annotation of the TR4 reference genome

Repeat masking identified repetitive sequences accounting for 5.02% of the curated TR4 nuclear genome. RNA-seq reads showed a 93.85% primary alignment rate and, together with fungal protein evidence, supported prediction of 8,898 protein -coding genes represented by 9,766 transcript models. Selection of one representative model per gene yielded 8,898 proteins containing 96.7% complete BUSCO orthologues, with only 0.4% duplicated BUSCOs. This representative set was used for downstream analyses; functional annotations are provided in Table S1.

### 3.3 Pathogenicity-related protein repertoires and secondary metabolic potential of TR4

Of the 8,898 representative TR4 proteins, 593 were classified as putatively secreted, including 258 candidate effectors (Figure 1A, B; Table S1). dbCAN3 identified 355 CAZyme candidates, including 137 in the putative secretome (Figure 1A, C), and fungiSMASH identified 20 candidate biosynthetic regions across seven nuclear contigs (Figure 1A, D).

**Figure 1.**
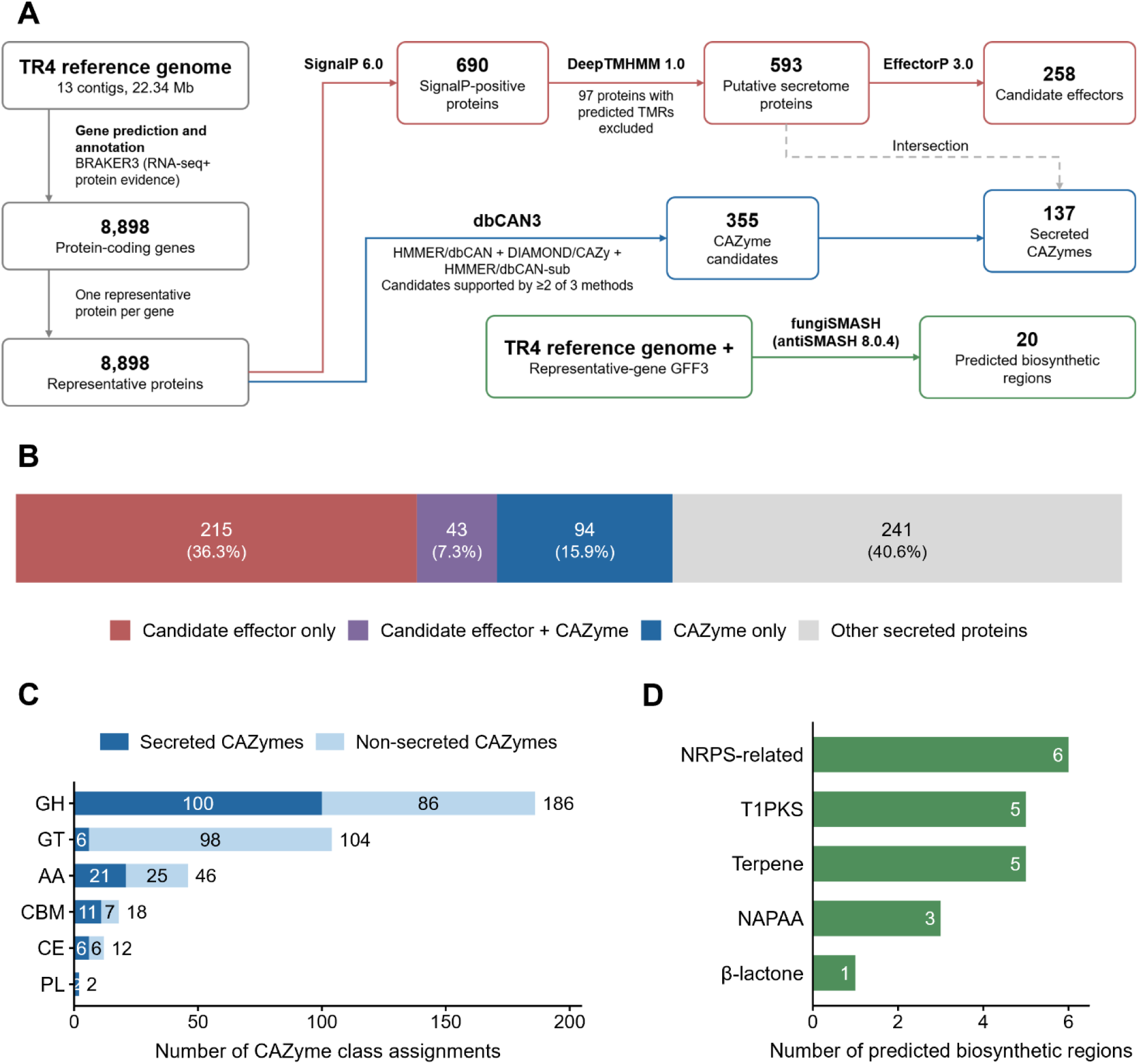
Functional genomic annotation and predicted secretome, carbohydrate-active enzyme (CAZyme) and biosynthetic repertoires of *Teratoramularia rumicicola* strain TR4. (A) Overview of the annotation and prediction workflow. From 8,898 representative proteins, 690 were SignalP-positive; exclusion of 97 proteins with predicted transmembrane regions yielded a putative secretome of 593 proteins, of which 258 were classified as candidate effectors. dbCAN3 identified 355 CAZyme candidates, including 137 in the putative secretome, and fungiSMASH identified 20 predicted biosynthetic regions. (B) Mutually exclusive composition of the 593 -member putative secretome according to candidate-effector and CAZyme classifications. (C) Distribution of CAZyme class assignments, separated into secreted and non-secreted CAZymes. CAZyme classes were not mutually exclusive. (D) Classification of the 20 predicted biosynthetic regions. GH, glycoside hydrolase; GT, glycosyltransferase; AA, auxiliary activity enzyme; CBM, carbohydrate - binding module; CE, carbohydrate esterase; PL, polysaccharide lyase; NAPAA, non-α-poly-amino acid; NRPS, nonribosomal peptide synthetase; T1PKS, type I polyketide synthase.

PHI-base-guided enrichment analysis identified significant enrichment of CAZymes and secreted CAZymes in both the whole-proteome and strict PHI-base-hit comparisons, with additional enrichment of the putative secretome and candidate effectors in the latter (Table S3). However, no category remained significant after multiple-testing correction when proteins whose only qualifying PHI-base phenotype was effector activity were reassigned to the comparison group. Among the 137 predicted secreted CAZymes, 45 had virulence-related PHI-base homologues. Of these, seven were prioritized as host-barrier-associated candidates based on predicted functions related to plant cell wall or cuticle modification (Table S4).

### 3.4 Genome-wide conservation and isolate-fixed sequence variation between TR4 and GR1

Pairwise whole-genome comparison showed that TR4 and GR1 were highly conserved, with 99.8084% average nucleotide identity. One-to-one alignments covered 21,106,518 bp of the TR4 assembly and 21,105,328 bp of the GR1 assembly, corresponding to 94.50% and 99.42% of their respective nuclear assembly spans.

Read-based comparison against the TR4 HiFi reference identified 28,040 high-confidence isolate-fixed differences, comprising 27,512 SNPs and 528 short indels (Figure 2A, B). Of these, 12,065 occurred within coding sequences. SnpEff classified 139 variants as HIGH impact and 6,513 as MODERATE impact, indicating that extensive genome-wide conservation coexisted with variation at a subset of coding loci with potential functional consequences.

**Figure 2.**
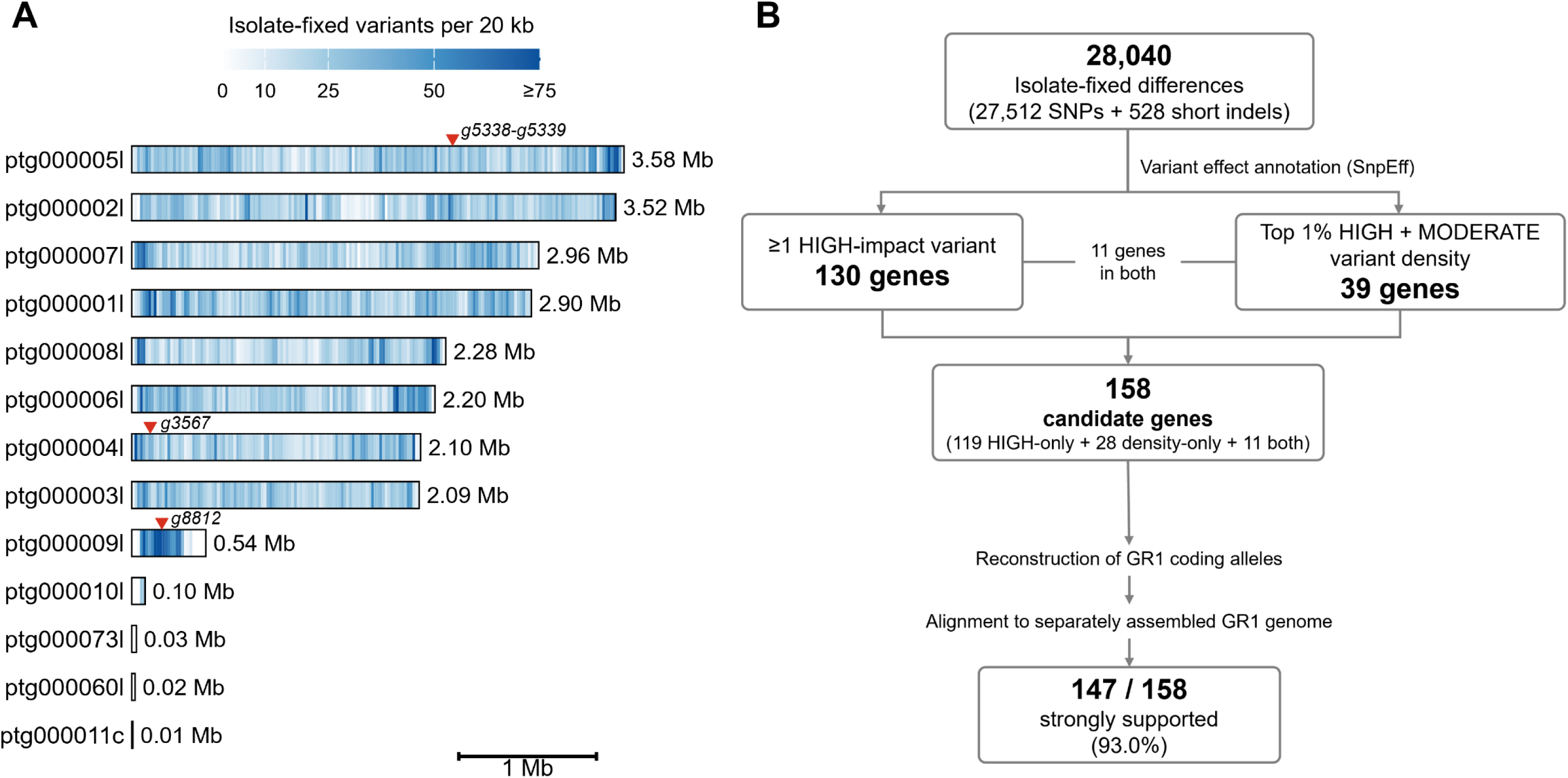
Genome-wide distribution of isolate-fixed sequence variation and systematic selection of candidate genes in *Teratoramularia rumicicola*. (A) Distribution of 28,040 high-confidence isolate-fixed differences between TR4 and GR1 across the 13 contigs of the TR4 HiFi reference genome, summarized in non-overlapping 20-kb windows. Window counts represent retained isolate-fixed differences and were not normalized by the number of callable positions. The positions of g5338 –g5339, g3567 and g8812 are indicated. (B) Workflow for systematic candidate-gene selection and assembly-based evaluation. Variant-based criteria identified 158 nonredundant candidate genes, of which 147 met the predefined criteria for strong support from the separately assembled GR1 genome.

### 3.5 Systematic reconstruction and assembly-based evaluation of candidate GR1 alleles

Application of the predefined criteria yielded 158 nonredundant candidate genes (Figure 2B; Table S2). Among the reconstructed GR1 alleles, 28 showed net coding - sequence length changes that disrupted the reading frame, eight lacked the annotated start codon and 55 contained a premature stop codon; these categories were not mutually exclusive. All 158 reconstructed coding sequences aligned to the separately assembled GR1 genome, and 147 met the criteria for strong assembly support. The remaining alleles showed minor sequence differences, partial coverage or mapping ambiguity. Separately, all seven prioritized host-barrier-associated secreted CAZyme genes retained intact coding sequences in GR1, with four identical proteins and three differing by only one or two amino-acid substitutions (Table S5).

### 3.6 Selected loci showing severe isolate-associated divergence

The clearest difference associated with a predicted biosynthetic region occurred at a nonribosomal peptide synthetase (NRPS)-like locus containing g5338 and the adjacent core biosynthetic gene g5339 (Figure 3A). The reconstructed GR1 allele of g5339 retained an intact 1,036 -amino-acid open reading frame with two predicted amino-acid substitutions (T522S and Q574K; Figure 3D). In contrast, a single -base insertion in g5338 caused a frameshift and premature termination, reducing the predicted Zn(II)₂Cys₆ transcription factor from 510 to 163 amino acids (Figure 3B, C).

**Figure 3.**
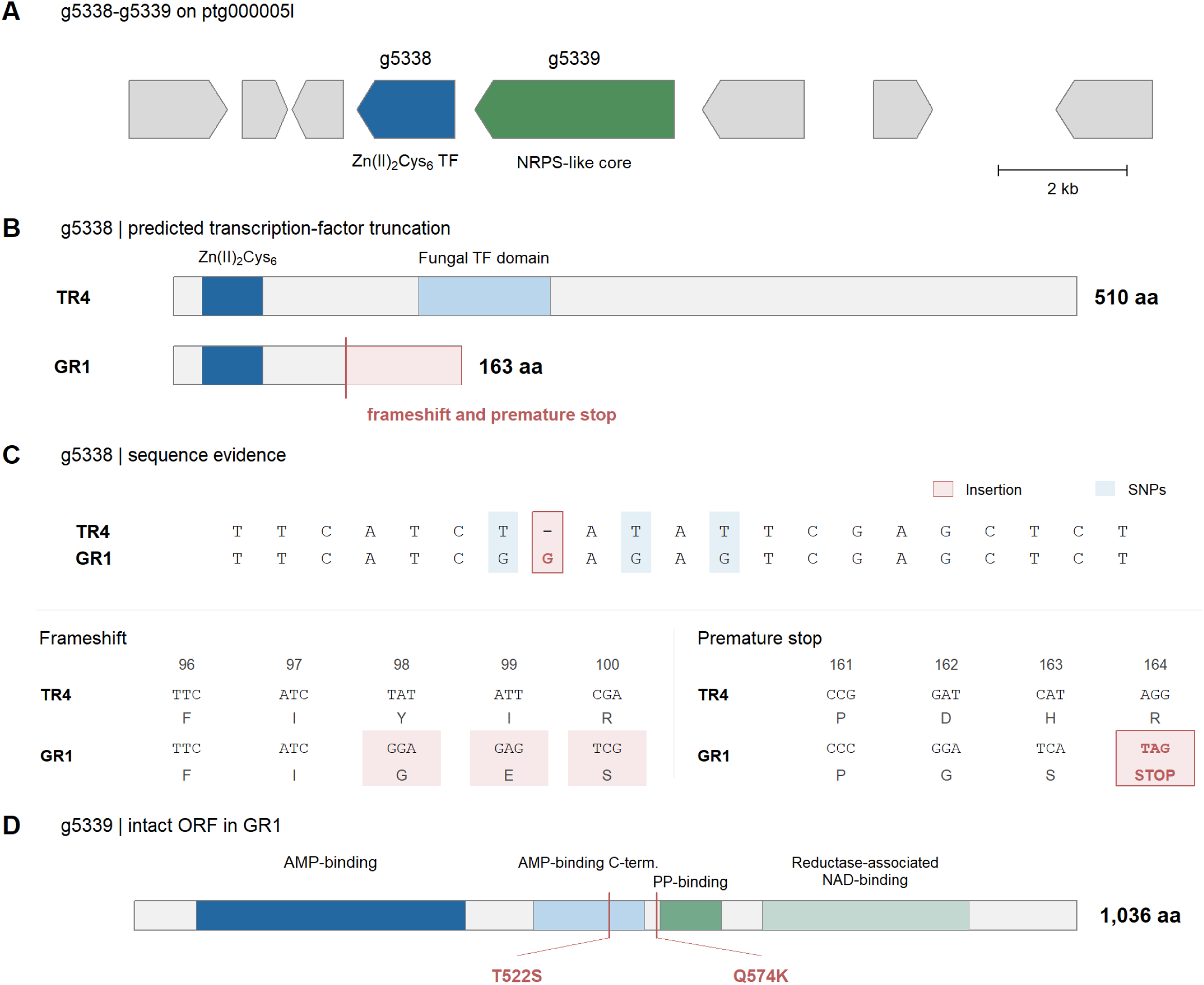
Predicted truncation of a Zn(II)_2_Cys_6_ transcription factor adjacent to a nonribosomal peptide synthetase (NRPS)-like core biosynthetic gene in *Teratoramularia rumicicola*. (A) Genomic organization of g5338 and g5339 on TR4 contig ptg000005l. Arrows represent annotated genes and indicate transcriptional orientation. (B) Predicted g5338 protein products in TR4 (510 amino acids) and reconstructed GR1 (163 amino acids), showing the frameshift-associated truncation. (C) Nucleotide and codon comparisons showing a single-base insertion and three nearby substitutions, the resulting frameshift at codon 98, and the first premature stop codon (TAG) at codon 164 in GR1. (D) Predicted domain organization of the TR4 g5339 protein and positions of two amino acid substitutions in GR1 (T522S and Q574K). The reconstructed GR1 allele retains an intact 1,036-amino-acid open reading frame. Domain annotations are based on the TR4 reference proteins and were not independently predicted for GR1. GR1 alleles were reconstructed assuming the TR4 exon and splice structures.

A second Zn(II)_2_Cys_6_ transcription-factor gene, g3567, was similarly disrupted by a two-base deletion, reducing the predicted protein from 558 to 21 amino acids (Table S2). At g8812, the reconstructed GR1 allele contained 42 fixed coding SNPs, including four premature-stop substitutions; the earliest yielded a predicted 53-amino-acid product compared with 186 amino acids in TR4 (Figure 4 A).

**Figure 4.**
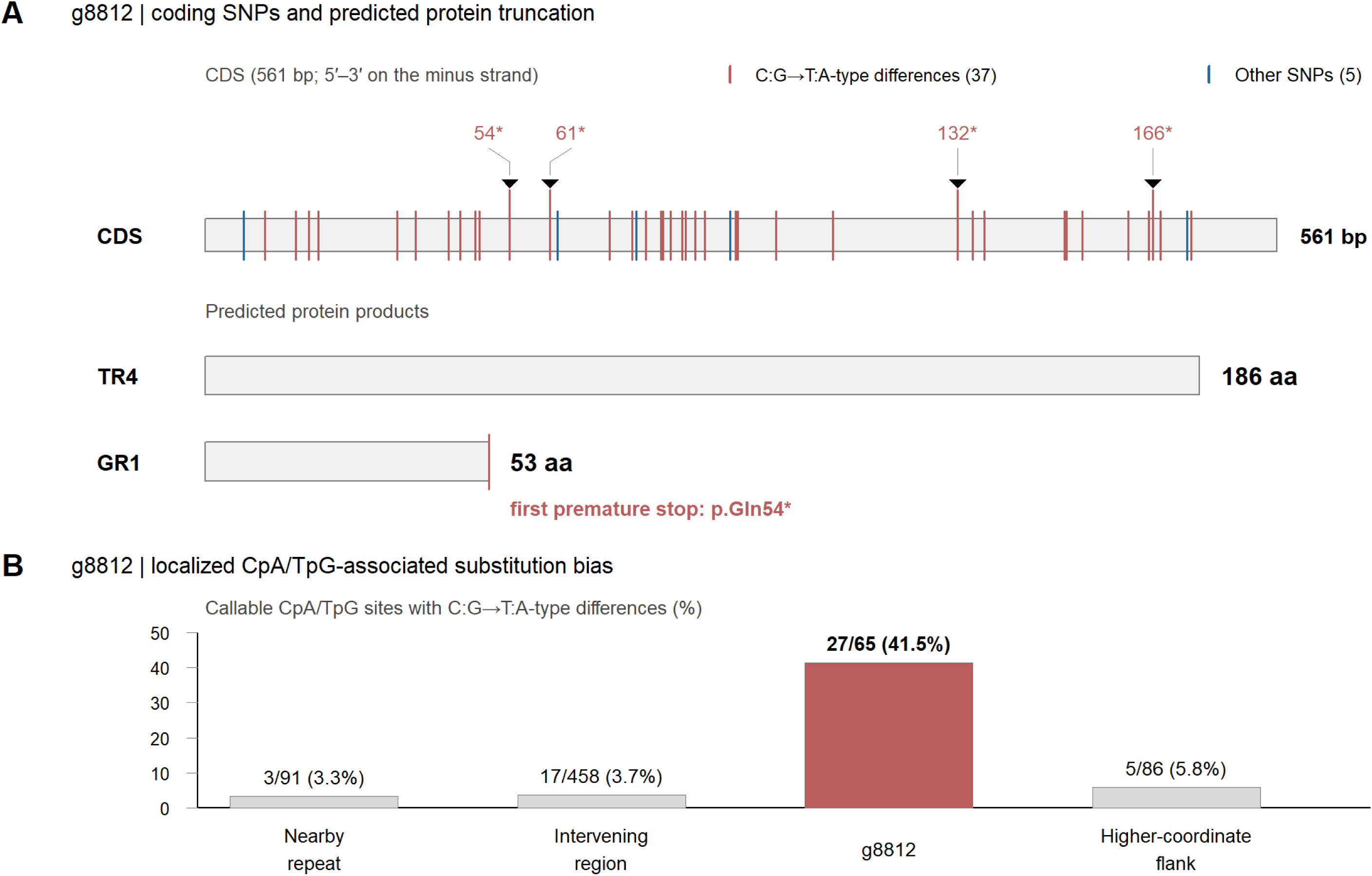
Coding-sequence disruption and localized RIP-like substitution patterns at the g8812 locus in *Teratoramularia rumicicola*. (A) Distribution of 42 isolate-fixed SNPs across the 561 -bp g8812 coding sequence (CDS), shown in the 5′–3′ orientation of the minus-strand gene. Red and blue vertical lines indicate 37 C:G-to-T:A differences and five other SNPs, respectively. Black arrowheads indicate four premature stop codons in the reconstructed GR1 allele. The first premature stop (p.Gln54*) is predicted to truncate the protein from 186 amino acids in TR4 to 53 amino acids in GR1. The CDS and predicted protein products are shown on separate scales. (B) Proportion of callable CpA/TpG sites carrying C:G-to-T:A isolate-fixed differences in four consecutive regions of TR4 reference contig ptg000009l: a nearby RepeatMasker-annotated repeat, the intervening region, the g8812 CDS and the higher-coordinate flank. The g8812 bar is highlighted in red, whereas the surrounding regions are shown in grey. Values above the bars indicate the number of mutated sites relative to callable CpA/TpG sites. Sites with read depths of 10–500 in both isolates were considered callable. Regions are arranged in increasing genomic-coordinate order.

The reconstructed g8812 CDS aligned uniquely across its full length to the GR1 assembly with 99.82% sequence agreement and therefore did not meet the ≥99.9% threshold for strong assembly support. Local read alignments nevertheless covered the entire g8812 CDS in both isolates, with mean depths of 205.6× in TR4 and 231.2× in GR1 and a mean mapping quality of 60 in both isolates after mapping-quality ≥30 and base-quality ≥20 filtering. Thirty-seven of the 42 substitutions were C:G-to-T:A changes, and the corresponding substitution frequency at callable CpA/TpG sites was 41.54% within g8812 versus 3.30 –5.81% in neighbouring regions (Figure 4A, B), consistent with a localized repeat-induced point mutation (RIP)-like substitution pattern.

Other candidates showed more limited predicted sequence changes despite meeting the selection criteria, underscoring the value of allele-level reconstruction (Table S2).

## 4. Discussion

This study provides a highly contiguous TR4 reference genome and the first structural and functional gene annotations for *T. rumicicola*, together with the first genome-scale comparison between isolates. Functional annotation yielded a catalogue of putative host-interaction factors, while comparison of TR4 and GR1 revealed extensive nucleotide conservation alongside pronounced predicted coding -sequence differences at selected loci (Figures 1 –4; Tables S1, S2).

The TR4 PacBio HiFi assembly substantially improved genome contiguity relative to the previously published short-read assembly, providing a highly contiguous reference genome for *T. rumicicola* (Table 1). The newly assembled GR1 genome extends genomic resources to a second isolate for the first time, establishing a basis for comparative genomic studies within the species. Furthermore, structural and functional gene annotations were generated for the first time in *T. rumicicola*, with structural gene prediction supported by RNA-seq and fungal protein evidence (Table S1).

Functional annotation of the TR4 genome provided a catalogue of putative host-interaction factors, including predicted secreted proteins, candidate effectors, CAZymes and secondary-metabolite biosynthetic regions (Figure 1; Table S1). Among these, CAZymes, particularly secreted CAZymes, showed significant enrichment for pathogenicity-related PHI-base homologues in both the whole-proteome and strict-hit analyses, although significance was lost in the sensitivity analysis excluding effector-activity-only phenotypes (Table S3). Seven secreted CAZymes were prioritised based on virulence-related PHI-base homology and predicted activities associated with plant cell wall or cuticle modification (Table S4). Their reconstructed GR1 alleles retained intact coding sequences with only limited amino-acid differences from TR4 (Table S5). However, their predicted functions and sequence conservation do not establish their involvement in pathogenicity.

This study provides the first genome-scale comparison between *T. rumicicola* isolates, revealing high nucleotide conservation between TR4 and GR1, with approximately 99.81% average nucleotide identity across aligned regions (Figure 2A). Nevertheless, read-based analysis identified 28,040 high-confidence isolate-fixed differences, comprising SNPs and short indels. These differences provided a basis for identifying candidate genes with potential coding consequences and examining locus-specific variation in subsequent analyses (Figures 2 –4; Table S2). However, the comparison was limited to two isolates, and its resolution was constrained by the fragmented GR1 assembly and the reliance on TR4 gene models for allele reconstruction.

Two Zn(II)₂Cys₆ transcription-factor genes, g5338 and g3567, contained frameshifts predicted to cause major protein truncation in GR1 (Figures 3B, C and Table S2). Zn(II)₂Cys₆ transcription factors can regulate secondary-metabolite biosynthesis in plant-pathogenic fungi, as demonstrated for SirZ in *Leptosphaeria maculans* (Fox et al., 2008). The proximity of g5338 to the intact NRPS-like core gene g5339 raises the possibility of altered secondary-metabolite regulation in GR1, although their regulatory relationship remains unverified. The predicted disruption of g3567 may also affect gene regulation and associated biological traits, but its targets and functions remain unknown.

Separately, g8812 showed a high density of C:G-to-T:A substitutions, with an elevated substitution frequency at CpA/TpG sites relative to flanking regions, and predicted premature termination (Figure 4A, B). RIP-associated sequence diversification has been reported in other plant-pathogenic fungi (Rouxel et al., 2011), suggesting that this unusual locus may warrant further investigation in the context of genome evolution. However, neither the ancestral state nor a duplicated target sequence was established, and the reconstructed g8812 allele showed reduced agreement with the GR1 assembly (Table S2). Together, these loci represent candidates for investigating potential molecular differences between isolates, although their functional and evolutionary significance remains to be determined.

Improving the efficacy of fungal microbial herbicides remains a major challenge for practical application (Gressel, 2024). The genomic resources established here, particularly the predicted host-interaction factors and isolate-associated coding variation, provide a basis for investigating the molecular mechanisms underlying pathogenicity in *T. rumicicola*. Future infection-stage expression analyses and targeted functional studies, combined with standardized comparisons of pathogenicity and weed-suppression activity, will be essential to determine the biological significance of the highlighted genes and genomic differences. Although genomic approaches may also contribute to production, formulation and biosafety, understanding pathogenicity mechanisms and isolate - associated variation could ultimately support strain selection, improvement and more effective use of *T. rumicicola* as a microbial herbicide.

## Acknowledgements

We thank Dr. Toyozo Sato (Niigata Agro-Food University, Niigata, Japan) for providing the *T. rumicicola* strains TR4 and GR1 used in this study. This work was supported by JSPS KAKENHI Grant Numbers JP25K23640 and JP24KJ1012.

## Author Contributions

M.I. conceived and coordinated the study, prepared biological samples, developed analytical workflows, performed data analyses, secured funding, prepared figures and drafted the manuscript. T.O. contributed to methodology, data analysis and validation. All authors critically reviewed and revised the manuscript and approved the final version.

## Data Availability Statement

The TR4 HiFi and GR1 nuclear genome assemblies generated in this study have been deposited at DDBJ/ENA/GenBank under accession numbers JBSLQR020000000 and JCERJE010000000, respectively. Raw sequencing reads are available under accessions SRR39794996 (TR4 PacBio HiFi), SRR39910844 (TR4 RNA-seq), and SRR39889942 (GR1 DNBSEQ). The previously published TR4 short -read assembly and raw sequencing reads are available under accessions GCA_054122135.1 and DRR802130, respectively. Supporting data are provided in the Supporting Information. The TR4 structural gene annotation and TR4 –GR1 isolate-fixed variant data are publicly available on Zenodo at https://doi.org/10.5281/zenodo.23333868.

## Artificial Intelligence Use Disclosure

ChatGPT (OpenAI) was used to assist with language refinement, manuscript drafting, the development and troubleshooting of computational workflows and analysis scripts, and the interpretation and visualisation of results. All AI-assisted content and outputs were reviewed and verified by the authors, who retained responsibility for the analyses, interpretations and final manuscript.

